# Ca_V_3.1 T-type calcium channels regulate spatial memory processing in the dorsal subiculum

**DOI:** 10.1101/2022.04.27.489701

**Authors:** Srdjan M. Joksimovic, Seyed Mohammadreza Ghodsi, Jasper A. Heinsbroek, James E. Orfila, Nicolas Busquet, Vesna Tesic, Robert Valdez, Brier Fine-Raquet, Vesna Jevtovic-Todorovic, Yogendra H. Raol, Paco S. Herson, Slobodan M. Todorovic

## Abstract

The dorsal subiculum (dSub) is one of the key structures responsible for the formation of hippocampal memory traces but the contribution of individual ionic currents to its cognitive function is not well studied. Although we recently reported that low-voltage-activated T-type calcium channels (T-channels) are crucial for the burst firing pattern regulation in the dSub pyramidal neurons, their potential role in learning and memory remains unclear. Here we used *in vivo* local field potential recordings and miniscope calcium imaging in freely behaving mice coupled with pharmacological and genetic tools to address this gap in knowledge. We show that the Ca_V_3.1 isoform of T-channels is critically involved in controlling neuronal activity in the dSub *in vivo*. Altering burst firing pattern by inhibiting T-channel activity markedly affects calcium dynamics, synaptic plasticity, neuronal oscillations and phase-amplitude coupling in the dSub, thereby disrupting spatial learning. These results provide a crucial causative link between the Ca_V_3.1 channels, burst firing activity of dSub neurons and memory processing, thus further supporting the notion that changes in neuronal excitability regulate memory trace formation. We posit that subicular Ca_V_3.1 T-channels could be a promising novel drug target for cognitive disorders.

## INTRODUCTION

Spatial navigation is a fundamental cognitive ability in both animals and humans that relies on intact function of the temporal lobe, particularly the hippocampal formation (Burgess et al., 2002). A growing body of evidence suggests an important role of the subiculum, the main output structure of the hippocampal formation, in processing of spatial memory traces (O’Mara, 2006; Aggleton and Christiansen, 2015). The subiculum receives information processed in the hippocampus through extensive unidirectional projections from the neighboring CA1 area (Amaral et al., 1995), along with relatively “unrefined” information directly from the cortical areas including the entorhinal cortex (O’Mara et al., 2009). Importantly, several recent studies identified a key role of the dorsal subiculum (dSub) neurons in contextual (Roy et al., 2017) and spatial information processing (Cembrowski et al., 2018).

Most subicular neurons are capable of generating burst firing (Jarsky et al., 2008; Joksimovic et al., 2017), a high-frequency barrage of action potentials, which is thought to be an important mechanism of distributing information across brain networks (Lisman, 1997; O’Mara et al., 2009). Burst firing may increase the fidelity of synaptic transmission and thereby modulate synaptic plasticity and neuronal signaling (Lisman, 1997; Izhikevich et al., 2003). Both *in vitro* and *in vivo* studies have shown that postsynaptic bursting activity is necessary for associative synaptic plasticity in the hippocampus, suggesting a crucial role of this firing pattern in learning and memory (Pike et al., 1999; Thomas et al., 1998). In line with these findings, Cembrowski et al. (2018) recently demonstrated that chemogenetic inhibition of bursting, but not regular-spiking neurons in the dSub leads to spatial learning deficits. However, the underlying mechanisms that drive excitability and burst firing of dSub neurons, as well as their functional implications related to mnemonic processing, are not well understood.

Due to their distinct biophysical properties and specific somatodendritic localization, low-voltage activated T-type calcium channels are ideally suited for the modulation of neuronal excitability and oscillatory activity (Iftinca and Zamponi, 2009). Specifically, the overlap of membrane potentials at which T-channels open, close, and inactivate allows for a basal inward flux of calcium ions, known as the ‘window’ current, which helps fine-tune the neuronal firing patterns (Williams et al., 1997). Another important property of T-channels is that their activation, even after a small membrane depolarization, may yield a low-threshold calcium spike crowned with a burst of action potentials. Our previous work using *ex vivo* slice preparation demonstrated that the Ca_V_3.1 isoform of T-channels is essential for controlling neuronal excitability of subicular neurons, particularly the bursting ones, as well as synaptic plasticity at CA1-subiculum synapses (Joksimovic et al., 2017). In the present study, we aimed to investigate whether this T-channel isoform, by supporting burst firing in dSub, may regulate neuronal oscillations *in vivo* and behavior associated with memory processing. To this end, we used a combination of miniscope calcium imaging, local field potential recordings and behavioral tasks for assessing spatial and contextual learning and memory. Our data indicate that pharmacological inhibition, global deletion of Ca_V_3.1 or dSub-specific molecular knockdown of Ca_V_3.1 T-channels reduces excitability of dSub neurons *in vivo*, alters local field potentials and cross-frequency phase-amplitude coupling of theta and gamma oscillations, which in turn produces deficits in memory processing. Our study for the first time reveals a crucial link between the T-channel-mediated burst firing in the dSub, neuronal oscillations *in vivo* and cognitive function in mice.

## MATERIAL AND METHODS

### Drugs

TTA-P2 (3,5-dichloro-N-[1-(2,2-dimethyl-tetrahydro-pyran-4-ylmethyl)-4-fluoropiperidin-4-yl methyl]-benzamide; Alomone Labs, Israel), a pan-selective T-channel blocker (Shipe et al., 2008), was dissolved/suspended in 15% 2-hydroxypropyl-β-cyclodextrin (Santa Cruz Biotechnology, Santa Cruz, CA, USA) immediately prior to *in vivo* calcium imaging experiment and administered intraperitoneally in a volume of 10 mL/kg.

### Animals

All procedures were conducted during the light cycle in male adult (2-5 month old) C57BL/6J (RRID:IMSR_JAX:000664) and Ca_V_3.1 knockout (KO; Ricken BioResources Centre, Japan) mice. Animals were housed in an AAALAC accredited animal facility according to protocols approved by the Institutional Animal Care and Use Committee of the University of Colorado Anschutz Medical Campus. Efforts were made to minimize animal distress and to use a minimal number of animals necessary to produce reliable scientific data. All experiments were conducted in compliance with the ARRIVE guidelines (Kilkenny et al., 2010).

### Virus delivery and GRIN lens implantation

C57BL/6J mice were anesthetized using a combination of 90-100 mg/kg ketamine and isoflurane (0.5-1% isoflurane, 2.5 L/min oxygen) and transferred to a motorized (NeuroStar, Germany) or standard stereotaxic frame (Kopf Instruments, CA, USA). Unilateral craniotomies 2-3 mm wide were created over the target region (AP= −2.80, ML= ±1.10) using a drill bit. To express GCaMP6f in principal dSub neurons, mice were stereotaxically injected with AAV1-CaMKIIa-GCaMP6f-WPRE.bGHppA (titer: 1.7 x 10^13^ vg/mL; Inscopix, Palo Alto, CA) using a 5 µL Hamilton syringe with beveled 31 G needle and 0.3 µL of the virus was delivered at DV= −1.90 at a rate of 0.1 µL/min. The syringe was retracted slowly and kept 200 µm above the injection site for 10 minutes to allow for diffusion of the virus. The same surgical procedure was performed in the Ca_V_3.1 T-channel short-hairpin RNA (shRNA) study in the dSub. Mice were first randomized into two treatment groups and then injected bilaterally into the dSub with 0.5 µL of scrambled control (AAV2-GFP-U6-scrmb-shRNA; titer: 1.1 x 10^13^ GC/mL; Vector Biolabs, Malvern, PA) or shRNA targeting Ca_V_3.1 (AAV2-GFP-U6-mCACNA1G-shRNA; titer: 6.8 x 10^12^ GC/mL; Vector Biolabs). Instead of a cranial window we drilled two small holes on each side of the skull at the appropriate coordinates. The shRNA target sequence was taken from Gangadharan et al. (2016). A subset of animals used only for *ex vivo* electrophysiological recordings received four injections into the dSub (anterior and posterior). Following virus infusions, the scalp was closed with sutures, and animals were allowed to recover for at least two weeks prior to experiments to ensure adequate virus expression.

The gradient index (GRIN) ProView™ Prism Lens probes (1.0 mm x 4.0 mm) with incorporated baseplates were purchased from Inscopix (part ID 1050-004637). After viral injection, the lens-incorporated baseplate was slowly lowered to the target region (AP= −2.80, ML= ±1.10, DV= 1.7) using the baseplate holder arm and stereotaxic frame following the manufacturer instructions. Three anchor-screws were implanted in the skull and the lens-incorporated baseplate was affixed to the skull with black dental cement to block light from the miniscope emission LED. All animals that underwent surgery were treated post-operatively with an analgesic (Banamine^®^, Merck) for two days. In the 4-8 week after surgery, mice were handled routinely by attaching the miniscope to the baseplate to get accustomed to the miniscope. Before the start of the experiment, a baseline measurement was recorded in an open-field arena (45 cm x 45 cm, black box).

### Calcium Imaging and Data Analysis

A counterbalanced within-subject design was used wherein mice were given an intraperitoneal (i.p.) injections of vehicle (15% cyclodextrin), 5 mg/kg TTA-P2 and 7.5 mg/kg TTA-P2 20 min prior to the recording sessions, and sessions for individual mice were separated by one week. Importantly, in this dose range, TTA-P2 is devoid of any sedation-like effects (Choe et al., 2011). During recording sessions, a miniscope (Inscopix nVoke) was mounted on the baseplate of mice with implanted GRIN lenses to record calcium activity in the dSub. Subjects were placed in the center of a novel environment (50 cm x 50 cm, white box), and allowed to freely explore the arena during imaging sessions. In each session, calcium activity was recorded in two 10 min epochs with a 10 min interval to reduce photobleaching.

Calcium imaging datasets were processed from raw miniscope videos using a specialized software (Data Processing Software, Inscopix). Videos were spatially and temporally downsampled by a factor of two, bandpass filtered, and movement corrected. Afterwards, a background subtraction (ΔF/F) was performed, and individual cells were identified using a Principal Component and Independent Component Analysis (PCA/ICA) algorithm. Identified cells were manually inspected for quality (high signal to noise calcium events in temporal components and cell-shaped non-overlapping spatial components). Calcium traces of selected cells were exported for the analysis of calcium event frequency and amplitude analyses. To track the activity of individual neurons across three imaging sessions, a longitudinal registration algorithm was applied to the spatial components identified by the PCA/ICA algorithm. Registered cells across imaging cells with a confidence threshold above 0.75 were selected for within-subject data analyses.

### Immunohistochemistry (IHC)

Brains of mice injected with Ca_V_3.1-shRNA or AAV-GCaMP6f, and implanted with a GRIN lens in dSub were processed for localization/placement verification. Mice were deeply anesthetized with 5% isoflurane and transcardially perfused with phosphate buffered saline (PBS pH 7.4, Life Technologies), followed by 4% paraformaldehyde in 0.1 M phosphate buffer, pH 7.4 (PFA). Whole brains were extracted and post-fixed in PFA for 24 h. Brains were rinsed in PBS, embedded in 3% agarose and brain sections (50 µm) were prepared on a microtome (Leica VT1200). Slices were rinsed three times in PBS, mounted on slides and antigen retrieval process was performed by exposing slides to a boiling citric buffer solution (0.1 M, pH 6.0). Sections were permeabilized in 1% glycine in PBST for 15 min and rinsed in PBS for 5 min. Sections were then blocked with 5% normal donkey serum in PBST (0.1% Triton X-100 in PBS) for 30 min, and incubated with primary antibody (rabbit anti-GFP; 1:1000; A11122; Invitrogen) diluted in 1% normal donkey serum in PBST overnight at 4 °C. Slices were rinsed 3 × 5 min in PBST followed by a 5 min rinse in PBS and then incubated for 2 h with secondary anti-rabbit antibody (anti-rabbit Alexa 488: 1:500, Invitrogen) at room temperature and washed 3 × 5 min in PBS. Sections were coverslipped using a fluorescent mounting medium containing DAPI (Vector laboratories) and images were taken using a confocal laser scanning microscope (Olympus FluoView FV1200) at 20× magnification using image stitching to obtain the entire region of interest. Only the mice that had viral GFP expression localized to the dSub were included in analysis.

### Electrode implantation, LFP recording and spectral analysis

To record local field potential (LFP)/electroencephalogram (EEG) signals, mice were implanted with two coated tungsten electrodes in the dSub [anterioposterior (AP)=-2.8 mm from bregma, mediolateral (ML)=±1.0 mm from midline, and dorsoventral (DV)=-1.9 mm below the skull surface) and two screw cortical EEG electrodes (AP=-1.0 mm, ML=±3.0 mm from midline) under ketamine (90-100 mg kg^-1^) and isoflurane anesthesia (0.5–2%). Screw electrodes placed behind the lambda on each side of the midline served as ground (right) and reference (left). The electrodes were fixed to the skull using dental acrylic. Lidocaine (1%) was injected at the surgery site to provide local anesthesia. The mice were treated post-operatively with an analgesic (Banamine^®^, Merck) for two days. At least one week after surgery, the synchronized, time-locked video and LFP/EEG signals were recorded from mice using the Pinnacle system (Pinnacle Technology Inc., Lawrence, KS). Acquired LFP/EEG signals were amplified 100 times and digitized at a sampling frequency rate of 2000 Hz (with a 0.5 Hz high-pass and 40 Hz low-pass filter) and stored on a hard disk for offline analysis. In the first experiment, WT and Ca_V_3.1 KO mice were put in a novel environment consisting of a Plexiglas cage specialized for EEG recordings with new bedding, food pellets and a water bottle. On the second day, a subgroup of animals was tested in a different environment (an alternate type of bedding with a set of new cues on the cage walls). After completion of experiments, mice were anesthetized with ketamine (100 mg/kg, i.p.) and electrolytic lesions were made by passing a 5 μA current for 1 s (5 times) through the depth electrodes. Mice were subsequently anesthetized with isoflurane and perfused with ice-cold 0.1 M phosphate buffer containing 1% potassium-ferrocyanide. Afterwards, brains were extracted, kept in 4% formalin (PFA) for 1-2 days and sliced (100–150 μm) using a vibratome (Leica VT 1200 S). Images of coronal slices with electrode locations were obtained using a bright-field Zeiss stereoscope and Zen Blue software (Zeiss, Germany). Some slices were counterstained with Nissl staining to help visualize and confirm electrode placements.

### Power spectral analysis

LFP/EEG recordings were taken during the first several hours after the animals was put in the novel environment, but the first 10 min and 60 min data points were analyzed using the power spectral analysis (fast Fourier transformation). The LFP frequency spectrum was subdivided in the following frequency bands: delta (0.5-4 Hz), theta (4-8 Hz), alpha (8-13 Hz), beta (13-30 Hz), and low gamma (30-40 Hz). The power spectral analysis (absolute power and power spectral density) and the spectrograms (Hann window size 2048, 50% overlap) were calculated using LabChart 8 software (ADInstruments Inc., Colorado Springs, CO). Although LFP signals from both right and left dSub were analyzed, only the recordings from the left dSub were used in the current study, as we obtained more reliable and artifact-free recordings from this side. Cortical EEG recordings are not presented in this study.

### Phase-amplitude coupling

Cross-frequency phase-amplitude coupling in LFP of the dSub was determined in MATLAB using phase-amplitude coupling (PAC) analysis (Cohen, 2014; J et al., 2015). For each mouse a continuous LFP time series of 1 h was down-sampled to 500 Hz and binned into 20 s epochs to increase signal-to-noise. Each epoch was flanked by 4 s buffer zones to prevent edge artifacts. Phase and power were determined by convolving binned data with 5-cycle complex Morlet wavelets. Phase was defined as the angle, and power defined as the squared magnitude of the convolved data. PAC was determined using the following formula:

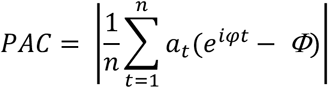

Wherein *t* denotes time, *a_t_* the power of the modulated frequency, *φ* the phase of the modulating frequency at *t*, and *i* the imaginary operator. The term *Φ* signifies a phase clustering debias term that was included to remove phase clustering artifacts that occur when field potentials (e.g. theta waves in the hippocampal formation), have non-sinusoidal characteristics (Belluscio et al., 2012)(J et al., 2015) This term is defined as:

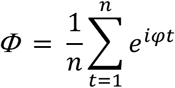

PAC values were normalized by z-transforming individually observed PAC values to a null-distribution generated by randomly shifting phase values relative to power values over 1000 iterations (PACz), and PAC at a given frequency pair was determined by averaging PACz values across all 20 s bins.

### Behavioral tests

For all behavioral experiments, the investigators were blinded to genotype and experimental treatment. Mice were subjected to a comprehensive behavioral characterization using tests aimed to assess motor, affective and cognitive functions, in the following order: open field, Y-maze, novel object/place recognition test, radial arm water maze, and/or contextual fear conditioning. Behavioral tests were separated by at least three days. All tests were performed by trained researchers and monitored and analyzed using specialized software (Ethovision system, Noldus, Wageningen, The Netherlands). Behavioral experiments in virus-injected mice began at least two weeks after injection to allow for adequate expression levels.

The open-field arena consisted of a grey polycarbonate box (44 cm × 44 cm × 30 cm). Mice were placed in the center of the arena, and tested for 10 min without prior habituation. This test provided important information about the general locomotor activity (total distance traveled), as well as anxiety-like behavior (number of entries and time spent in the center of the arena) of mice.

The Y-maze consists of a three-arm maze with equal angles between all arms (30 cm long and 5 cm wide with 12 cm high walls). Mice were initially placed in the center of the arena, and the sequence of arm entries was recorded manually by a trained observer for each mouse over an 8-min period, in order to quantify alternations.

In the novel object recognition test, an individual adult mouse was placed into an opened-top testing arena (30 cm x 30 cm square with 25 cm high walls), and allowed to investigate two unique objects for 5 minutes before being returned to its home cage. Ten minutes later, the mouse was again placed in the test arena and allowed to investigate one familiar (from previous exposure) and one novel object for 3 minutes. The time spent investigating the familiar and novel objects was recorded using a digital video camera interfaced with a computer. Specialized movement tracking software recorded the number and duration of visits to either object in the test arena.

Novel place recognition is the variation of the novel object recognition task, which focuses mostly on hippocampal memory. For 5 minutes, mice were exposed to 2 identical copies of an object in two corners of the arena. In the test trial (10 min delay), one of these objects was repositioned in a previously empty corner. The novel location of that familiar object should induce longer investigation.

The radial arm water maze (RAWM) apparatus consists of a circular pool (100 cm in diameter and 35 cm in height) with inserts forming six arms 60 degrees apart. One of these arms features a platform submerged 1 cm below the water surface. The room-temperature water was made opaque by adding a nontoxic water-soluble white dye, which also helps enhance the contrast between the mouse and the background for optimal tracking. The animals were tested for 2 consecutive days (15 swim trials per day, in 3 sessions of 6, 6 and 3 trials). Each swim trial lasted a maximum of 60 s, and unsuccessful mice were guided to the platform, in order to facilitate learning about the existence of the escape platform. After each trial, mice were gently dried with towels, put to a warm enclosure for 5-10 minutes, and then returned to their home cages at the end of each session. To assess spatial learning and memory, we quantified the latency to reach the platform and the number of errors (reference and working memory), respectively. For the Ca_V_3.1-shRNA experiment, the RAWM task was made more difficult by changing the position of the hidden platform on Day 2.

In the contextual fear conditioning paradigm, mice are first exposed to an electric shock in a specific context, and subsequent exposure to the same context (without the shock) should elicit fear-induced freezing behavior, indicating that the mice have formed an association between the context and the aversive stimulus. Mice were placed into conditioning boxes that are 30.5 cm x 24.1 cm x 29.2 cm in size. The context of the box included a metal grid floor, a house light, and an odor cue (70% Ethanol, which is also used for cleaning the apparatus). On the first day of training, the mice were placed into the chamber for 8 minutes during which 3 shocks (2 s, 0.7 mA) were presented. On the second day, mice were placed into the same context as day 1 (same box, but no shock applied), and the mouse behavior was monitored for 8 minutes, but only the first 3 minutes were analyzed for freezing behavior. Mice were followed up to 72 h, by placing them again in the same context.

### Brain slice preparation for patch-clamp electrophysiology experiments

Patch-clamp experiments using brain slices from the virus-injected mice began at least two weeks after injection. Animals were anesthetized briefly with isoflurane, decapitated, and their brains rapidly removed. Fresh horizontal brain slices, 250-300 µm thick, were sectioned at 4 °C in pre-chilled solution containing (in mM): sucrose 260, D-glucose 10, NaHCO_3_ 26, NaH_2_PO_4_ 1.25, KCl 3, CaCl_2_ 2, MgCl_2_ 2, using a vibrating micro slicer (Leica VT 1200S). Brain slices were immediately incubated for 45 minutes in a solution containing (in mM): NaCl 124, D-glucose 10, NaHCO_3_ 26, NaH_2_PO_4_ 1.25, KCl 4, CaCl_2_ 2, MgCl_2_ 2 at 37 °C prior to use in electrophysiology experiments, which were conducted at room temperature. During incubation, slices were constantly perfused with a gas mixture of 95 % O_2_ and 5 % CO_2_ (v/v).

### Patch-clamp electrophysiology recordings

The external solution for whole-cell recordings consisted of (in mM): NaCl 125, D-glucose 25, NaHCO_3_ 25, NaH_2_PO_4_ 1.25, KCl 2.5, MgCl_2_ 1, CaCl_2_ 2. This solution was equilibrated with a mixture of 95 % O_2_ and 5 % CO_2_ (v/v) for at least 30 minutes with a resulting pH of approximately 7.4. The internal solution for recording well isolated T-currents consisted of (in mM): tetramethyl ammonium (TMA)-OH 135, EGTA 10, MgCl_2_ 2, and Hepes 40, titrated to pH 7.2 with hydrofluoric acid (HF) (Todorovic and Lingle, 1998). For cell-attached and current-clamp experiments, the internal solution consisted of (in mM): potassium-D-gluconate 130, EGTA 5, NaCl 4, CaCl_2_ 0.5, HEPES 10, Mg ATP 2, Tris GTP 0.5, pH 7.2. Glass micropipettes (Sutter Instruments O.D. 1.5 mm) were pulled using a Sutter Instruments Model P-1000 and fabricated to maintain an initial resistance of 3-5 MΩ. GFP-expressing dSub neurons were identified using the microscope with epifluorescence and IR-DIC optics. T-channel activation was measured by stepping the membrane potential from a 3.6 s-long pre-pulse to the test voltage (from −120 to −50 mV). The single unit extracellular (loose) or cell-attached recordings were obtained at the holding membrane potential of 0 mV (Nunemaker et al., 2003). Intrinsic excitability of dSub neurons was characterized by using a multi-step protocol which consisted of injecting a family of depolarizing (50-150 pA) current pulses of 400 ms duration in 25 pA increments followed by a series of hyperpolarizing currents of the same duration stepping from −200 to −400 pA in 50 pA increments. Regular-spiking and depolarization-induced burst firing properties of dSub neurons were characterized by examining the membrane responses to depolarizing current injections at the membrane potential of −60±1 mV. All recordings were made in the presence of GABA_A_ and ionotropic glutamate receptor blockers [(20 μM picrotoxin, 50 μM D-2-amino-5-phosphonovalerate (D-APV) and 5 μM 2,3-dihydroxy-6-nitro-7-sulfamoyl-benzo[*f*]quinoxaline-2,3-dione (NBQX)] in the external solution. Subsequent action potential frequencies (per pulse and per burst) and input resistances were determined. The membrane potential was measured at the beginning of each recording and was not corrected for the liquid junction potential, which was around 10 mV in our experiments. The membrane input resistance was calculated by dividing the end of steady-state hyperpolarizing voltage deflection by the injected current. Neuronal membrane responses were recorded using a Multiclamp 700B amplifier (Molecular Devices, CA, USA). Voltage current commands and digitization of the resulting voltages and currents were performed with Clampex 8.3 software (Molecular Devices), and voltage and current traces were analyzed using Clampfit 10.5 (Molecular Devices). Spontaneous action potentials recorded in the cell-attached configuration were analyzed offline with MiniAnalysis (Synaptosoft, Inc., Decatur, GA, USA).

### Long-term potentiation

Mice (Ca_V_3.1-shRNA or scrambled control) were transcardially perfused with ice-cold (2–5 °C) oxygenated (95% O_2_/5% CO_2_) artificial cerebral spinal fluid (aCSF) for 2 min prior to decapitation. The composition of aCSF was the following (in mM): 126 NaCl, 2.5 KCl, 25 NaHCO_3_, 3 NaH_2_PO_4_, 2.5 CaCl_2_, 1.2 MgCl_2_ and 12 glucose. Hippocampal slices (300 μm) containing the dSub were dissected and transferred to a holding chamber containing aCSF for at least 1 h before recording. Field excitatory postsynaptic potentials (fEPSPs) were monitored using a half-maximal stimulus based on a baseline input–output curve. After a 20-min stable baseline was established, LTP was induced in the dCA1–dSub pathway by delivering a theta burst stimulation (TBS) train (four pulses delivered at 100 Hz in 30 ms bursts, repeated 10 times with 200 ms interburst intervals), as described previously (Toledo et al., 2017). The amount of potentiation was calculated as the percentage change from baseline (the averaged 10 min slope value from 50 to 60 min post-TBS divided by the averaged slope value at 10 min prior to TBS).

### Data Analysis and Statistics

The sample size was estimated using G*Power software (Faul et al., 2007) based on our previous experience (α = 0.05; power ≥ 0.8; effect size=0.8). Results are presented as mean ± standard deviation in the text and mean ± standard error of the mean in the graphical presentations. All data are biological replicates. Statistical analyses were performed using unpaired two-tailed t-tests, ordinary one-way or repeated measures analysis of variance (RM ANOVA), two-way RM or mixed-effects ANOVA (treatment or genotype as one factor, and time or current-injection as the other factor). Where applicable, Sidak’s or Tukey’s *post hoc* test for multiple comparisons was also used, as recommended by GraphPad Prism 9.2 software (GraphPad Software, La Jolla, CA). Fisher’s exact test was used to compare the proportions of different neuronal types in the dSub. Cohen’s d was calculated to measure effect size for the most substantive findings. Effect significance was accepted as P<0.05. Statistical and graphical analyses were performed using GraphPad Prism 9.2 and Origin 2018 (OriginLab, Northhampton, MA).

## RESULTS

### Pharmacological inhibition of T-channels decreases neuronal excitability in the dSub of freely behaving mice

We have showed an important contribution of T-channels to the activity of principal pyramidal neurons in the dSub using acute brain slices *ex vivo* (Joksimovic et al., 2017), but the role of these channels in excitability of intact dSub of freely moving mice is not known. To record from principal dSub neurons *in vivo*, mice were stereotaxically injected into the dSub with AAV carrying the calcium indicator GCaMP6f under control of the CaMKIIα promoter followed by the implantation of a 1-mm diameter GRIN lens tangent to the dSub (Figure 1A). *In vivo* calcium imaging sessions were performed at baseline and 20 min after i.p. injections of vehicle or TTA-P2 by placing the mouse in a novel open-field arena (Figure 1B). Successful GCaMP6f transfection and lens implantation were assessed by IHC in each subject 10-12 weeks post-surgery (Figure 1C). During each imaging session, calcium transients of neuronal populations were recorded in two epochs (first and last 10 min of a 30 minute session) and analyzed to derive the calcium transients. Figure 1D shows the calcium signals from five registered neurons across three recording sessions, whereas panels E and F depict the average calcium transient frequency and amplitude for all neurons within the recorded population. The dose-dependent decrease in calcium transients’ amplitude after i.p. injection of relatively low doses of TTA-P2 demonstrates the critical role of T-type calcium currents in neuronal excitability of dSub. Specifically, we observed a 20% decrease in the average calcium amplitude after the injection of 5 mg/kg TTA-P2 (F_2,371_=33.79, P<0.001; P=0.004), and about 35% decrease after 7.5 mg/kg TTA-P2 (P<0.001; Cohen’s d=1.20), as compared to the vehicle (Figure 1E). Importantly, the calcium activity recorded after vehicle injection was not different from activity during the baseline measurement. The inhibitory effects of TTA-P2 were limited to the amplitude of calcium transients, and not the frequency (Figure 1F). We did note a slightly higher frequency after 7.5 mg/kg, as compared to the lower dose (F_2,371_=8.42, P<0.001; P<0.001), but this could be ascribed to a larger number of calcium transients of smaller amplitude that are revealed after a more pronounced T-channel antagonism. These findings were confirmed by the cumulative probability distribution plots of calcium amplitude (Figure 1G) and the inter-event intervals (Figure 1H): a leftward shift after TTA-P2 was evident for the amplitude, but not the frequency of calcium transients. To examine this data in more detail, we tracked the activity of a subset of individual dSub neurons across different imaging sessions, and we confirmed that the addition of TTA-P2 significantly decreased the calcium amplitude (Figure 1–figure supplement 1; F_2,84_=79.88, P<0.001; P=0.028 for 5 mg/kg and P<0.001 for 7.5 mg/kg vs. vehicle; one-way RM ANOVA followed by Tukey’s *post hoc* test). These *in vivo* calcium imaging data sets confirm our prior *ex vivo* results, thus providing further support for the notion that Ca_V_3.1 T-channels are important regulators of dSub neuronal activity.

**Figure 1.**
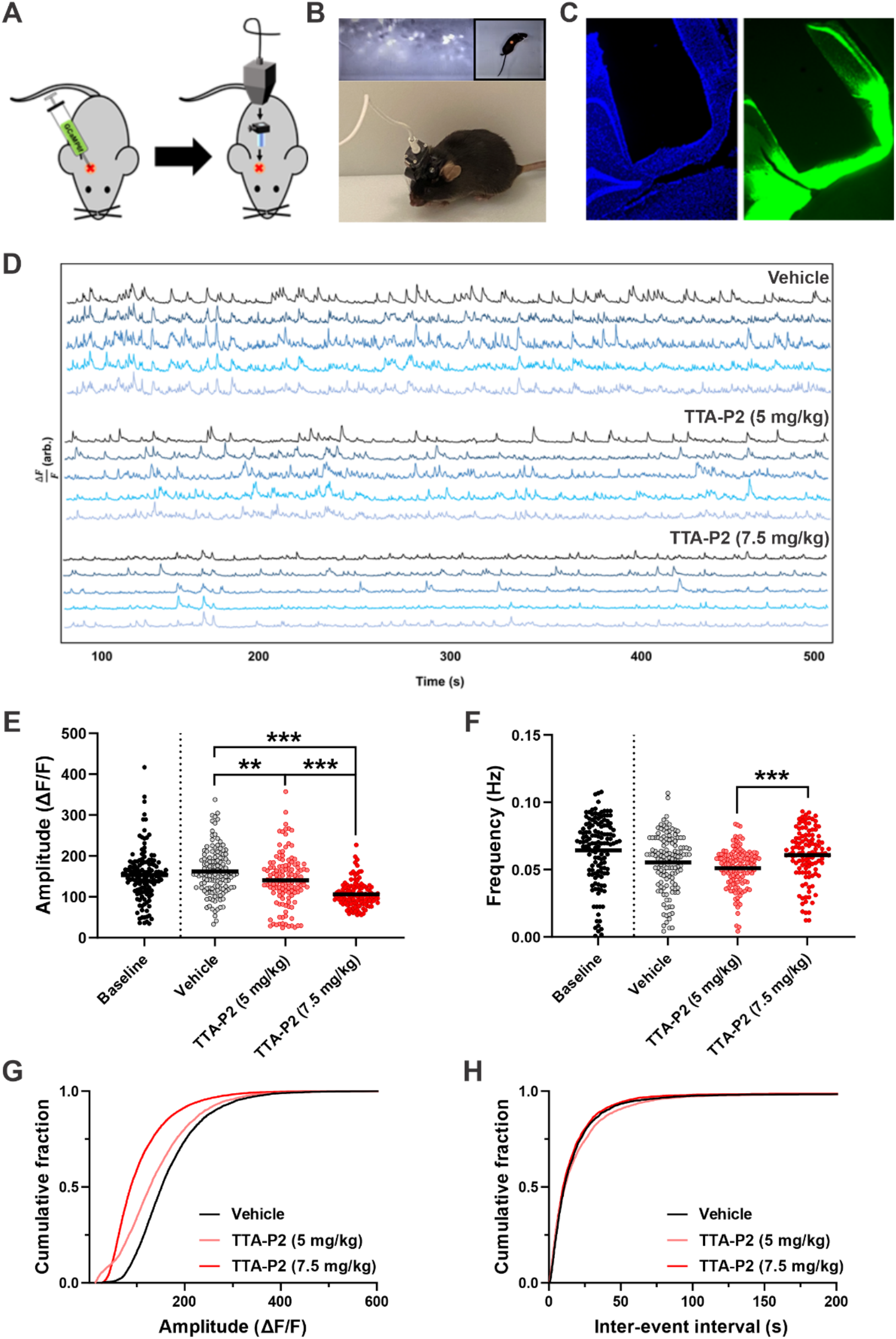
The effects of selective T-channel antagonist (TTA-P2) on dSub neuronal activity in freely behaving mice recorded using *in vivo* calcium imaging. **A.** The cartoon depicts the implantation procedure of the miniscope, which starts with the injection of an AAV for the viral transduction of principal dSub neurons with GCaMP6f (AAV1-CaMKII-GCaMP6f) followed by the GRIN lens and baseplate implantation. **B.** An example imaging field of GCaMP6f-expressing dSub neurons captured with the miniscope camera, is presented along with the actual photo of a mouse with the mounted miniscope in the open-field arena. **C.** Representative coronal section showing GCaMP6f-labeled neurons (green) and the lens placement right above the dSub; counter-stained with DAPI (blue). **D.** Original calcium traces recorded from five representative dSub neurons after the i.p. injection of vehicle (top), 5 mg/kg TTA-P2 (middle) and 7.5 mg/kg TTA-P2 (bottom) in mice exploring the open-field arena. **E.** The average amplitude of calcium transients recorded over four different sessions: baseline (black; n=134 neurons, 4 mice), vehicle (gray; n=130 neurons, 4 mice), 5 mg/kg TTA-P2 (pink; n=128 neurons, 4 mice) and 7.5 mg/kg TTA-P2 (red; n=116 neurons, 3 mice). **P<0.01 and ***P<0.001, one-way ANOVA followed by Tukey’s *post hoc* test. **F.** The average frequency of calcium transients recorded during the same four sessions. **G.** The cumulative probability plots of the calcium amplitude after the injection of vehicle (8687 events), 5 mg/kg TTA-P2 (7896 events) or 7.5 mg/kg TTA-P2 (8838 events). **H.** The cumulative probability plots of the inter-event intervals after the injection of vehicle, 5 mg/kg TTA-P2 or 7.5 mg/kg TTA-P2. **P<0.01 and ***P<0.001, one-way ANOVA followed by Tukey’s *post hoc* test.

### Ca_V_3.1 T-channels regulate neuronal oscillations and phase-amplitude coupling in the mouse dSub during exploration of a novel environment

Changes in neuronal excitability may profoundly affect oscillatory activity in the hippocampus, particularly in the gamma range (Klemz et al., 2022). Based on this notion, and the fact that all hippocampal neurons are strongly theta-modulated (Buzsaki et al., 2002), we hypothesized that decreasing neuronal excitability, including bursting, should hamper the generation of these oscillations in the dSub and their coupling to the theta wave, a phenomenon heavily involved in cognitive processing (Ole Jensen and Colgin, 2007). Since hippocampal low gamma oscillations in mice are important for the acquisition of spatial information in a novel context (Kitanishi et al., 2015; Trimper et al., 2017), we first examined the spectral power across a range of frequencies (≤ 40 Hz) in the dSub of WT and Ca_V_3.1 KO mice during the active exploration of a novel environment (Figure 2). The original traces and time-frequency domain heat maps from representative LFP recordings are presented in Figure 2 (panels A and B, respectively). We found reached statistical significance for beta (genotype: F_1,13_=7.62, P=0.016) and low gamma (genotype: F_1,13_=5.69, P=0.033) frequency bands. For example, the average low gamma power was decreased by about 70% in Ca_V_3.1 KO mice (Cohen’s d=1.13), as compared to the wild-type group. We noticed a similar pattern when we plotted the data using the power spectral density (PSD; Figure 2D), but only for the low gamma frequency range (genotype: F_1,13_=5.59, P=0.034). The PSD analysis also revealed that the spectral power of the 8-Hz theta wave was significantly affected in Ca_V_3.1 KO mice (interaction: F_4,52_=3.09, P=0.023; *post hoc*: P<0.001). Collectively, our results indicate that Ca_V_3.1 T-channels play a crucial role in regulating the dSub oscillatory activity predominantly in a beta-low gamma frequency range.

**Figure 2.**
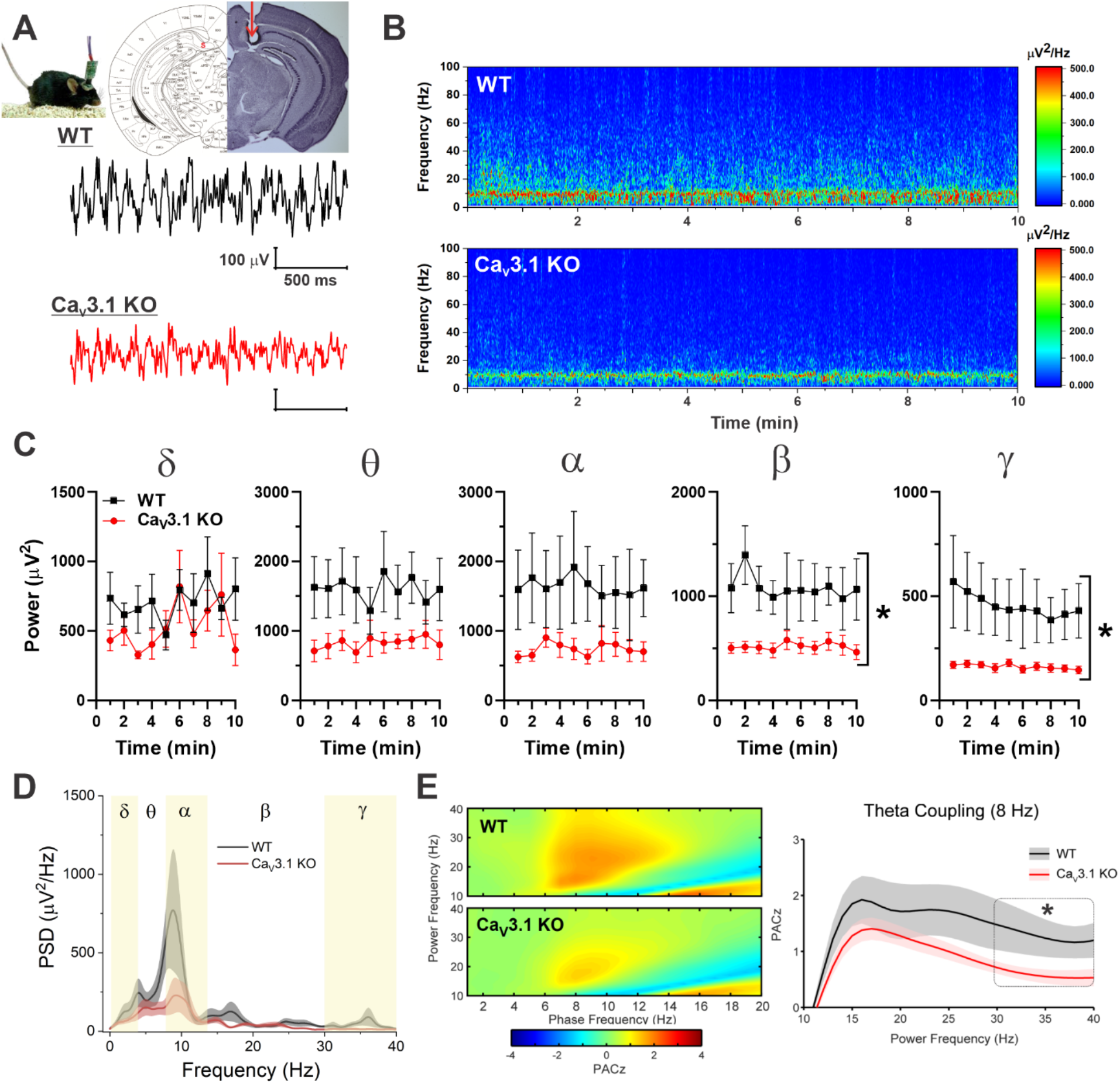
LFP recorded from the dSub of WT and Ca_V_3.1 KO mice during exploration of a novel environment. **A.** Top: Nissl staining of coronal brain slice with red arrow showing the electrode placement in the dSub. Bottom: Representative raw traces of subicular LFP from WT (black) and Ca_V_3.1 KO (red) mice. **B.** Representative spectrograms computed from the LFP recordings in WT (top) and Ca_V_3.1 KO (bottom) mice. Note the decrease in spectral power across different frequencies in Ca_V_3.1 KO mice. **C.** Time course plots show the changes in the absolute spectral power in different frequency bands in WT (n=6) and Ca_V_3.1 KO (n=9) mice while exploring a novel environment. *P<0.05, two-way RM ANOVA. **D.** The power spectral density (PSD) plot during the same 10-min time period in WT (black) and Ca_V_3.1 KO (red) mice. **E.** Contour plots depicting PACz values in WT and Ca_V_3.1 KO across a range of frequencies for both amplitude and phase. Note a difference between WT and Ca_V_3.1 KO mice in PAC between theta/alpha and beta/gamma bands. The PACz plots show the difference in alpha/beta/low gamma to theta (8 Hz) coupling between WT (n=6) and Ca_V_3.1 KO (n=9) mice exploring a novel environment. *P<0.05, two-way RM ANOVA.

Next, we investigated the effects of global Ca_V_3.1 deletion on phase-amplitude coupling (PAC) in the dSub. The coupling of the power of gamma oscillations to the phase of theta waves is a well-described phenomenon in the hippocampal formation linked to cognitive function (O. Jensen and Colgin, 2007); the amplitude of a higher frequency should vary depending on the phase of a lower frequency. In contour plots of averaged PAC data across a range of frequencies, Ca_V_3.1 KO mice displayed a significant reduction in the coupling of beta and gamma frequencies to theta and alpha waves compared to the WT group (Figure 2E, left panel). We next quantified the specific coupling of the high frequency oscillations to the theta wave (8 Hz), the best characterized driver frequency for power in higher frequency bands in the hippocampal formation, and we observed a pronounced decrease only in theta-low gamma coupling (Figure 2E, right panel; genotype: F_1,13_=5.02, P=0.043). Taken together, our dSub LFP data obtained from mice that are exploring novel environment demonstrate an important role of Ca_V_3.1 T-channels in regulating neuronal oscillations in this region, as well as maintaining adequate coupling between theta and higher frequencies, which may also impact cognitive behavior.

### Ca_V_3.1 KO mice show deficits in spatial learning

The behavioral phenotype of Ca_V_3.1 KO mice is not fully characterized. Although deficits in novelty seeking have been reported in these mice (Gangadharan et al., 2016), the function of the hippocampal formation has not been assessed in detail. Based on our LFP findings presented above, as well as our published long-term potentiation (LTP) data (Joksimovic et al., 2017), we reasoned that global Ca_V_3.1 deletion may have important behavioral consequences related to hippocampal-dependent learning and memory. The open-field test capitalizes on the conflicted motivation of rodents to investigate new environment, yet avoid exposed situations. Therefore, we first evaluated the general locomotor activity of these mice, and we found no significant difference compared to WT mice, as assessed by the total distance traveled in the open-field arena (Figure 3A; 5262±674 cm in the WT group vs. 4671±968 cm in the KO group). In addition, these mice displayed no marked differences in the time and the number of entries to the central zone of the arena, the two anxiety-related parameters (Figure 3, panels B and C, respectively). Next, we investigated whether Ca_V_3.1 KO mice show deficits in gross hippocampal function by assessing the spontaneous alternation behavior in the Y-maze (Figure 3D). The Y-maze is a relatively simple, non-invasive, test for measuring spatial working memory, which heavily depends on the hippocampal activity (Hughes, 2004). We did not detect any short-term spatial memory deficits in this test (66.0±9.1% in the WT group vs. 67.0±11.8% in the KO group), which was in line with our premise that the overall hippocampal processing is not severely affected in these mice. To further explore the functioning of the hippocampal formation in Ca_V_3.1 KO mice, we tested these two group of mice in the novel place and novel object exploration tests (Figure 3E). Novel place recognition test focuses mostly on hippocampal memory, whereas the novel object recognition depends mostly on perirhinal cortex, and is a highly utilized memory testing procedure that takes advantage of rodents’ natural preference to investigate objects they have not previously encountered; therefore, mice tend to spend more time with novel than familiar objects (Dere et al., 2007; Barker et al., 2007). In both paradigms, Ca_V_3.1 KO mice were indistinguishable from figure supplement 1A; t_16_=0.68, P=0.504) or novel object (Figure 3E; t_16_=0.90, P=0.380). One sample t-test revealed that WT mice, unlike Ca_V_3.1 KO mice, preferred investigating a novel object significantly above the 50% chance level (t_9_=2.67, P=0.026 and t_7_=0.53, P=0.611, respectively), suggesting that certain deficits in hippocampal-perirhinal activity may exist in Ca_V_3.1 KO mice. Finally, we tested these two cohorts of mice in a radial arm water maze (RAWM), a relatively challenging but robust paradigm that can detect nuanced changes in spatial learning and memory in mice (Alamed et al., 2006). Using this paradigm, we observed a significant impairment in spatial learning in Ca_V_3.1 KO mice, as evidenced by longer latencies to reach a hidden platform in the water maze across sessions compared to WT controls (Figure 3F; genotype: F_1,23_=4.74, P=0.040). The total number of the memory errors, however, was not significantly different from the control group (Figure 3–figure supplement 1B; genotype: F_1,23_=1.77, P=0.197). Our behavioral data suggests that the spatial information processing is selectively disrupted in mice lacking Ca_V_3.1 T-channels.

**Figure 3.**
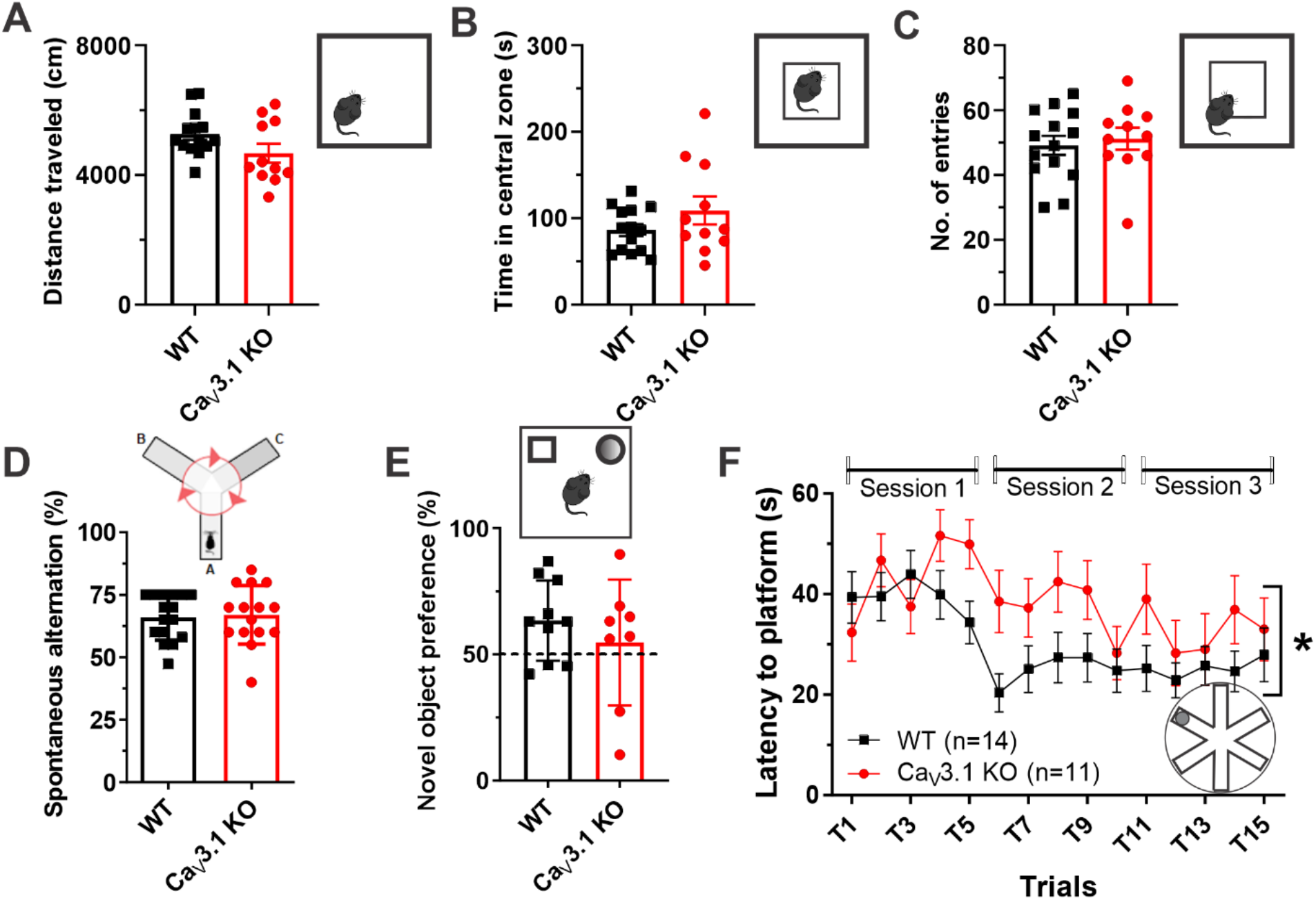
Behavioral assessment of general activity, anxiety-related behavior and cognitive functions in Ca_V_3.1 KO mice. **A.** The distance traveled of WT (n=14) and Ca_V_3.1 KO (n=11) mice in the open-field arena. **B.** The time spent in the central zone of the open-field arena. **C.** The number of entries to the central zone of the open-field arena. **D.** The percentage of spontaneous alternation of WT (n=16) and Ca_V_3.1 KO (n=15) mice in the Y-maze test. **E.** The preference in the time exploring a novel object in WT (n=10) and Ca_V_3.1 KO (n=8) mice during the novel object recognition test. **F.** The latency to reach the hidden platform in the RAWM in WT (black) and Ca_V_3.1 KO (red) mice. *P<0.05, two-way RM ANOVA.

### Selective tissue-specific silencing of Ca_V_3.1 gene attenuates burst firing in the dSub

Our results obtained with global knockout mice are important to validate the contribution of Ca_V_3.1 T-channels to different hippocampal-dependent behaviors. However, this T-channel isoform is found throughout the cortex and other parts of the hippocampal formation (Talley et al., 1999). Therefore, we next utilized a more selective shRNA-based viral approach for the bilateral knockdown (KD) of Ca_V_3.1 expression in the dSub (Figure 4A). To confirm functional knockdown, we used patch-clamp recordings in acute brain slices to examine T-current properties in shRNA-expressing (GFP-positive) and neighboring (GFP-negative) neurons following shRNA or control AAV injections. The original traces of T-currents in representative dSub neurons are presented in Figure 4B. We identified a profound decrease of the T-current amplitude in GFP+ neurons in the Ca_V_3.1-shRNA group of about 67%, as compared to the control group (Figure 4C; interaction: F_1,23_=4.81, P=0.039; *post hoc*: P=0.003). Importantly, T-currents in GFP-neurons in these two treatment groups were of similar amplitude (Control: 269.9 ± 123.1 pA; Ca_V_3.1-shRNA: 244.4 ± 65.5 pA).

**Figure 4.**
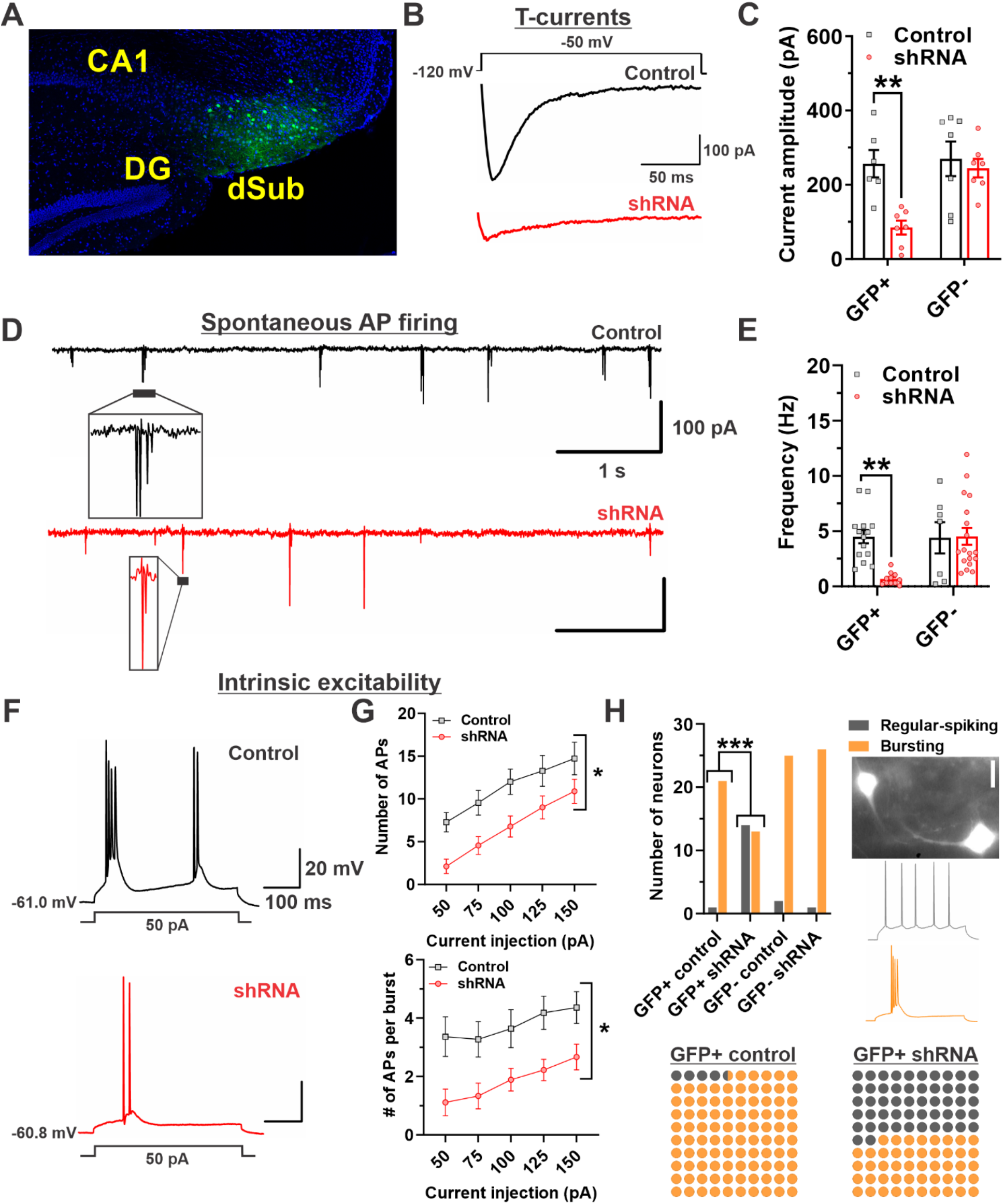
Electrophysiological properties of subicular neurons in dSub-specific Ca_V_3.1 knockdown mice. **A.** Representative image shows GFP-transfected neurons only in dSub (green); counter-stained with DAPI (blue). **B.** Original T-current traces from representative dSub neurons in mice injected with either scrambled shRNA (Control; black) or Ca_V_3.1-shRNA (shRNA; red), generated using a double-pulse protocol with a 3.6-s-long prepulse (from −120 mV to −50 mV). **P<0.01, two-way ANOVA followed by Sidak’s *post hoc* test. **C.** The average T-current amplitudes in GFP+ and GFP- neurons after control and shRNA viral injections (Control: n=6 in GFP+ and n=7 in GFP-; shRNA: n=7 per group). **D.** Original traces from representative GFP+ dSub neurons depicting cell-attached recordings of spontaneous action potentials (AP) in control (black) and shRNA-treated neurons (red). The inset shows a typical burst firing pattern, which appears attenuated after knockdown of Ca_V_3.1 channels. **E.** The average firing AP frequencies in GFP+ and GFP- neurons after control and shRNA viral injections (Control: n=14 in GFP+ and n=7 in GFP- groups; shRNA: n=11 in GFP+ and n=18 in GFP- groups). **P<0.01, two-way ANOVA followed by Sidak’s *post hoc* test. **F.** Original traces from representative dSub neurons showing the burst firing pattern in response to a depolarizing 50 pA current injection in control (black) and shRNA-treated neurons (red). **G.** The average number of APs per pulse (top) and the average number of APs per burst (bottom) recorded after a series of escalating current injections in dSub neurons transduced with control or Ca_V_3.1-shRNA. *P<0.05, two-way RM ANOVA. **H.** The number of GFP+ and GFP- regular-spiking and burst firing neurons after control and shRNA viral injections [GFP+ control: n=22 neurons (1 regular-spiking vs. 21 burst firing); GFP+ shRNA: n=27 (14 vs. 13); GFP- control: n=27 (2 vs. 25); GFP- shRNA: n=27 (1 vs. 26)]. Note that shRNA- treated mice have a large proportion of GFP+ regular-spiking neurons compared to control mice. The image shows two GFP-labeled dSub neurons. Original traces of a typical regular-spiking (top, gray) and burst firing neuron (bottom, orange) after a depolarizing 100 pA current injection. ***P<0.001, Fisher’s exact test.

Next, we recorded spontaneous action potential firing in a cell-attached (loose) configuration, and the original representative traces of GFP+ control (black) and Ca_V_3.1-shRNA (red) dSub neurons are presented in Figure 4D. We detected a significant decrease in AP frequency of GFP+ neurons in the Ca_V_3.1-shRNA group (Figure 4E; interaction: F_1,46_=5.99, P=0.018; *post hoc*: P=0.002 vs. Control GFP+ neurons), with burst firing being clearly diminished in the subset of tested neurons (see inset). Importantly, spontaneous AP firing was not different in non-virally transduced GFP-negative control neurons (Control: 4.4±3.8 Hz; Ca_V_3.1-shRNA: 4.5±3.2 Hz). Finally, we concluded our patch-clamp study by investigating the intrinsic excitability of dSub Ca_V_3.1 KD neurons. Similarly to our previous data in global Ca_V_3.1 KO mice (Joksimovic et al., 2017), the target gene silencing significantly affected the burst firing pattern of subicular neurons, as shown by representative traces in Figure 4F. Both the number of APs per current injection (Figure 4G top; treatment: F_1,18_=6.23, P=0.023) and per burst (Figure 4G top; F_1,18_=8.11, P=0.011) were significantly reduced in the Ca_V_3.1 shRNA group, as compared to the control virus group. Considering that the distal parts of the dSub contain predominantly burst firing neurons (Cembrowski et al., 2018; Jarsky et al., 2008), we detected an unusually high number of regular-spiking GFP+ neurons in dSub-specific Ca_V_3.1 KD mice (Figure 4H; 51.9% vs. 4.6%, P<0.001), which may indicate that a large portion of dSub neurons in this group switched their modes of firing from bursting to regular-spiking. In line with this finding, we previously reported that mice with global Ca_V_3.1 deletion have a lower fraction of burst firing neurons than wild-type mice (Joksimovic et al., 2017). Overall, our results show that the chosen AAV-shRNA approach successfully silenced the target gene in dSub neurons, which significantly affected neuronal excitability in this region, particularly the burst firing pattern. These data extend our previous findings in global Cav3.1 KO mice, and further confirm the importance of Ca_V_3.1 T-channels for the dSub neuronal activity.

### dSub-specific Ca_V_3.1 KD produces deficits in synaptic plasticity and spatial learning in mice

We previously reported that mice with the global deletion of Ca_V_3.1 T-channels show impaired synaptic plasticity at the CA1-Sub synapse (Joksimovic et al., 2017). However, it was not possible to deduce whether the observed deficit was caused entirely by changes in the dSub activity. Furthermore, this study was conducted using horizontal hippocampal slices that mostly contain intermediate and ventral portions of the hippocampal formation. To address these concerns, we used our AAV-shRNA Ca_V_3.1 KD approach to study LTP at the CA1-Sub synapse in the dorsal portions of the hippocampal formation using theta-burst stimulation, a protocol developed to mimic plasticity produced by theta-modulated gamma rhythms that occurs during learning and memory formation (Larson and Munkácsy, 2015). In line with our hypothesis that Ca_V_3.1-mediated excitability of dSub neurons is required for synaptic plasticity and learning and memory processing, we observed an impaired LTP at the dCA1-dSub synapse measured 60-70 minutes following TBS (Figure 5A; t_9_=2.679, P=0.025; Cohen’s d=1.55).

**Figure 5.**
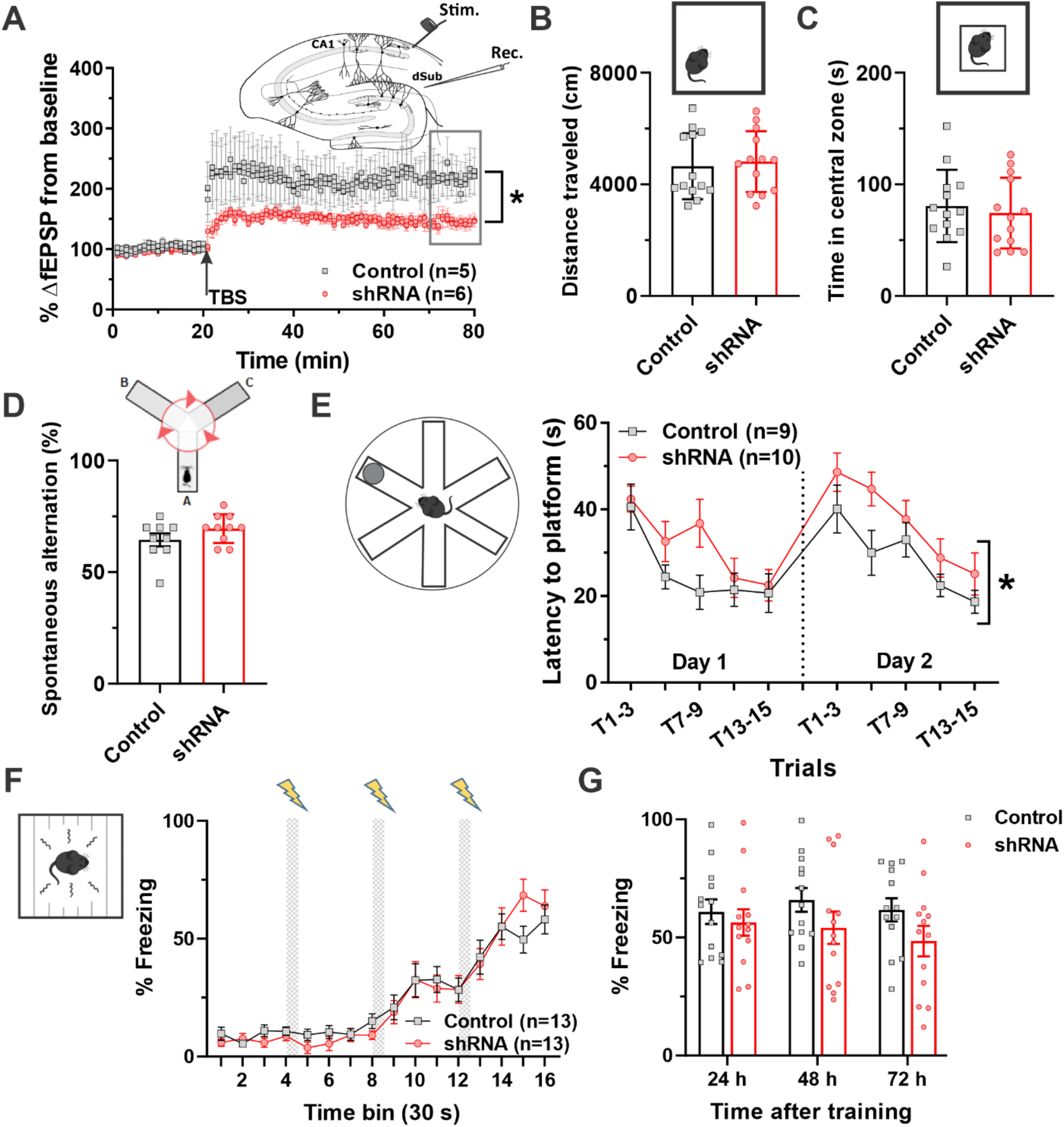
Synaptic plasticity and behavioral assessment of general activity, anxiety and spatial and contextual memory in dSub-specific Ca_V_3.1 knockdown mice. **A.** The time course of normalized fEPSP slope recordings at the dCA1-dSub synapse in control (gray squares) and Ca_V_3.1-shRNA (pink circles) mice. The inset shows a schematic diagram of hippocampal formation: stimulation probe for LTP induction in the dSub was located towards the end of CA1 in the stratum oriens/alveus region, whereas the recording pipette was in the molecular layer of the dSub. *P<0.05, t-test **B.** The distance traveled of control and Ca_V_3.1-shRNA mice in the open-field arena (n=13 mice per group). **C.** The time spent in the central zone of the open-field arena. **D.** The percentage of spontaneous alternation of control (n=9) and Ca_V_3.1-shRNA (n=10) mice in the Y-maze test. **E.** The latency to reach the hidden platform in the RAWM in control (black) and Ca_V_3.1-shRNA (red) mice. *P<0.05, two-way RM ANOVA. **F.** The percentage of time spent freezing in control (gray squares) and Ca_V_3.1-shRNA (pink circles) mice during the training phase of the contextual fear conditioning paradigm. **G.** The percentage of time freezing in control and Ca_V_3.1-shRNA mice during recall tests up to 72 h after contextual fear conditioning.

Next, we set out to investigate the behavioral effects produced by Ca_V_3.1 KD in the dSub that may arise from impaired synaptic plasticity as evidenced by impaired LTP and reduced excitability in dSub neurons. As expected based on our results obtained with global Ca_V_3.1 KO mice, silencing subicular Ca_V_3.1 T-channels did not affect general locomotor activity or anxiety-related behavior in an open-field (Figure 5, panels B and C, respectively). The values for the total distance traveled and the time spent in the central zone in both groups of injected mice were comparable to the wild-type control group presented in Figure 3 (panels A and B), which may indicate that Ca_V_3.1-shRNA, but also the entire stereotaxic injection protocol, did not alter motor functions, anxiety-related and novelty-seeking behavior in mice. Additionally, the spontaneous alternation behavior in the Y-maze was not affected by the dSub-specific Ca_V_3.1 KD procedure (Figure 5D; 64.4±8.8% in the control group vs. 69.5±6.4% in the Ca_V_3.1-shRNA group), suggesting that the spatial working memory does not rely on Ca_V_3.1 T-channel activity in the dSub.

Since we demonstrated selective inhibition of burst activity and LTP in the dSub with our Ca_V_3.1-shRNA approach, we reasoned that Ca_V_3.1 KD mice may show impaired spatial learning in the RAWM. Indeed, mice expressing Ca_V_3.1-shRNA in dSub neurons showed a spatial learning impairment, as evidenced by longer latencies to reach the hidden platform in the maze compared to control mice (Figure 5E; treatment: F_1,34_=5.63, P=0.024). Although relatively consistent, this effect was more pronounced on the second day of testing, when the platform position was changed, which may be associated with a decreased cognitive flexibility in these mice that is caused by deficits in acquisition of new and/or updating old spatial information, and not retrieval of previously acquired ones. To confirm this finding, we focused on the first three trials during the second day of testing and analyzed the rate of revisits to the arm that contained the hidden platform on the previous day, and we found no difference between the two groups (data not shown; t_17_=1.66, P=0.115), which suggests that Ca_V_3.1 KD mice did not show any deficits in retrieving the spatial memory trace. Furthermore, the total number of memory errors was also not significantly affected in Ca_V_3.1 KD mice (Figure 5–figure supplement 1; treatment: F_1,34_=2.66, P=0.112). Conversely, Roy et al. (2017) showed that optogenetic inhibition of dSub leads to deficits in retrieval of hippocampal-dependent contextual information. To clarify these two seemingly opposite findings, we examined whether decreasing activity of burst firing neurons in the dSub will impair fear memory retrieval using the contextual fear conditioning paradigm. The contextual fear conditioning is a hippocampal-dependent task that tests the association between a nociceptive stimulus and the context in which it occurred (Chen et al., 1996). Over the course of training, both dSub Cav3.1 KD and control mice showed a similar and expected behavioral response to the foot shock (Figure 5F). For example, during conditioning after the third foot shock, control mice spent 58.2±22.2%, and Ca_V_3.1 KD mice spent 63.8±24.5% of the time in freezing behavior. When the animals were returned to the same boxes for testing 24 hours later, no difference in their freezing behavior was observed (Figure 5G; Control: 60.9±18.8%; Ca_V_3.1-shRNA: 56.4±20.1%), indicating successful retrieval of the contextual fear memory; similar observations were made 48 and 72 hours after the training occurred (treatment: F_1,24_=2.05, P=0.165). Taken together, these results suggest that the spatial, but not fear memory formation is impaired in mice with silenced Ca_V_3.1 T-channel expression in the dSub.

## DISCUSSION

Alterations in synaptic plasticity and neuronal oscillations of the subiculum are critically involved in cognitive deficits related to a wide range of brain disorders (Godsil et al., 2013; Goutagny et al., 2013). Since these processes are regulated by neuronal excitability, the tendency of subicular pyramidal neurons to burst in rhythmic patterns is likely to be critical for processing cognitive information (O’Mara et al., 2009). However, ionic currents that regulate this specific neuronal activity are not well studied in the subiculum. Our previous study showed that Ca_V_3.1 is the most abundant T-channel isoform in the subiculum, and that it plays a major role in the regulation of neuronal excitability and synaptic plasticity in this part of the hippocampal formation (Joksimovic et al., 2017). Here, using a more targeted approach coupled with multiple *in vivo* techniques in freely behaving mice, we extend our findings to show that Ca_V_3.1 T-channels regulate calcium dynamics of dSub neurons, neuronal oscillations and cross-frequency coupling associated with information processing in the hippocampal formation. Furthermore, we show that global and dSub-specific silencing of Ca_V_3.1 function result in attenuated burst firing pattern, impaired synaptic plasticity, and spatial memory deficits. Collectively these data show a crucial role for Ca_V_3.1 T-channels in the functioning of dSub neurons within the larger hippocampal formation.

Our *in vivo* miniscope calcium imaging data clearly demonstrate that T-channels are functionally involved in maintaining somatic calcium levels and modulating activity of principal dSub neurons. The selective T-channel antagonist TTA-P2 (Shipe et al., 2008) produced a dose-dependent decrease in the amplitude of calcium transients recorded from dSub neurons in freely behaving mice. Previous work from our lab and others indicates that TTA-P2 essentially abolished burst firing in subicular (Joksimovic et al., 2017) and thalamocortical (Dreyfus et al., 2010) principal neurons in acute brain slices, which may imply that the observed decrease in calcium amplitude is associated with alterations in this firing pattern. Furthermore, a growing body of evidence suggests that it is possible to infer spiking activity from calcium imaging (Sasaki et al., 2008; Theis et al., 2016), and that bursts of action potentials are associated with an increased calcium transient amplitude, when compared to single spikes (Chen et al., 2013). These observations combined with our previous results in acute brain slices suggest that Ca_V_3.1 T-channels are the main driving force behind the bursting activity of dSub neurons in both *in vivo* and *ex vivo* conditions.

The subiculum generates gamma oscillations in a broad frequency range (30-150 Hz) (Jackson et al., 2011), which is known to facilitate neuronal communication and regulate the timing and topography of hippocampal output (Colgin and Moser, 2010). Interestingly, Eller et al. (2015) showed that bursting, but not regular-spiking neurons fire during gamma oscillations, which implies that modulating ionic currents that underlie bursting may affect the generation of these oscillations in the dSub. In humans, several studies showed that the oscillatory activity in the gamma range is associated with the formation of episodic memories (Axmacher et al., 2006; Sederberg et al., 2007). In rodents, slow hippocampal gamma oscillations are implicated in both memory acquisition (Trimper et al., 2017) and retrieval (Colgin, 2016), and provide an indication of synchronicity between different areas within the hippocampal formation (Colgin et al., 2009). We found that mice lacking Ca_V_3.1 T-channels display a broad-band decrease in spectral power across different frequencies in the dSub, but mid- to high- frequencies (beta/low gamma range) are particularly affected. Considering that gamma oscillations are required for effective information flow within the hippocampal formation, we propose that altering the generation of these oscillations in dSub would disrupt spatial memory processing in Ca_V_3.1 KO mice. In line with this hypothesis, we observed a diminished phase-amplitude coupling between theta and gamma oscillations in the dSub of these mice. This finding could be relevant to several brain disorders, Alzheimer’s disease in particular. For example, Goutagny et al. (2013) detected robust changes in theta-gamma coupling in the subiculum even before amyloid beta accumulation. Interestingly, a reduced expression of Ca_V_3.1 T-channels in the hippocampus was found in aged brains, which was exacerbated in patients with Alzheimer’s disease (Rice et al., 2014). Theta waves are the main regulatory mechanism for synchronization of information processing and synaptic plasticity in the hippocampal formation (Buzsáki, 2002), whereas theta-gamma coupling is a robust predictor of successful long-term synaptic potentiation and memory formation (Bikbaev and Manahan-Vaughan, 2008). Indeed, the neighboring CA1 principal neurons show phase-locked firing during low gamma oscillations in a novel environment, which is associated with strengthening of synaptic plasticity and the process of acquiring new information (Kitanishi et al., 2015). The only other study investigating the role of Ca_V_3.1 T-channels in hippocampal oscillations found that theta power was increased in the medial septum, which was accompanied by the hyperexcitability of septo-hippocampal GABA-ergic neurons in mice lacking this T-channel isoform (Gangadharan et al., 2016). As we previously reported, T-channels are only modestly involved in modulating firing properties of subicular GABAergic interneurons (Joksimovic et al., 2017), thus we believe that the decrease in neuronal excitability of pyramidal excitatory neurons, predominantly the bursting ones, caused the observed changes in oscillatory activity. This notion is supported by our calcium imaging and Ca_V_3.1-shRNA viral experiments, which selectively targeted principal dSub neurons. Our study provides novel insights into the role of T-channels in regulating oscillatory activity of intact hippocampal networks.

The proximal part of the dSub (adjacent to CA1 area) is enriched with regular-spiking neurons that project to the nucleus accumbens and medial prefrontal cortex, to name a few, whereas the distal part contains almost exclusively burst firing neurons that primarily receive inputs and project back to the medial entorhinal cortex (Cembrowski et al., 2018), a brain structure crucial for spatial navigation (Tukker et al., 2021). Based on the fact that bursting is an important and reliable means of transferring information (Lisman, 1997), and bearing in mind their specific projection pattern, one could presume that inhibiting burst firing neurons would have specific consequences on memory formation. In line with this idea, a selective chemogenetic inhibition of the distal, but not proximal, part of the dSub is associated with deficits in acquisition of spatial information, rather than retrieval (Cembrowski et al., 2018). Our results from the RAWM largely confirm this finding, as dSub-specific KD mice exhibit deficits in spatial learning but do not appear to have any deficit in recollecting where the hidden platform was on the previous day or have any reference memory impairments. We also demonstrated an impairment in dCA1–dSub synaptic plasticity in these mice following theta-burst stimulation, which further supports this hypothesis. The data from the contextual fear conditioning revealed no difference between dSub-specific Ca_V_3.1 KD and control of mice suggesting that the acquisition and immediate retrieval of fear memory traces were not strongly dependent on activity of burst firing neurons in dSub. Using optogenetics, Roy et al. (2017) showed that inhibiting dSub projections to medial entorhinal cortex impairs context-dependent memory updating, a process important for retrieval of contextual information. Having said this, our AAV-shRNA approach will inherently have more subtle effects on dSub neuronal activity than optogenetics, and it mostly affects neurons in the distal dSub that project to several brain areas, which may explain these seemingly opposite findings. Taken together, our LFP, LTP, and behavioral data indicate that Ca_V_3.1 T-channels in the dSub are particularly important for the formation of spatial memory traces.

Several studies showed that the dSub contains spatially tuned place cells (Sharp, 1997; Sharp, 2006) and vector-based cells (Lever et al., 2009), which indicates its involvement in the spatial code processing within the hippocampal formation. Using high-resolution juxtacellular recordings in freely moving rats, Simonnet and Brecht (2019) reported that bursting neurons contribute more to spatial information processing than regular-spiking dSub neurons, demonstrating a more important role of this firing pattern in subicular space coding. Along with lower intrinsic excitability of burst firing dSub neurons, we observed a smaller proportion of these neurons in mice with altered Ca_V_3.1 T-channel expression, which would inherently affect spatial information processing and consequently impair spatial navigation; our behavioral results in the RAWM using both global Ca_V_3.1 KO and dSub-specific Ca_V_3.1 KD mice provide strong support for this hypothesis.

It is well established that alternating between different modes of firing generates oscillatory patterns in the thalamocortical circuitry, which are important for normal sensory processing, attention, sleep-wake transitions, and consciousness (Llinás et al., 2005; Steriade et al., 1991). Thus, dysfunction of T-channels may lead to different pathophysiologies, including epilepsy (Khosravani and Zamponi, 2006), changes in arousal and altered sleep-wake patterns (Anderson et al., 2005). The role of the Ca_V_3.1 isoform in mediating these behavioral abnormalities is well documented (Kim et al., 2001; Lee et al., 2004), but little is known about its contribution to the anxiety-like behavior, learning and memory, and cognition. Our behavioral results in both global Ca_V_3.1 KO and dSub-specific Ca_V_3.1 KD mice reveal a new role of these channels in spatial information processing. Importantly, both of these mouse lines did not exert any substantial changes in general motor activity, anxiety-like behavior or gross hippocampal function, which validates our findings related to cognitive behavior.

In conclusion, we report that Ca_V_3.1 T-type calcium channels are the essential regulators of neuronal excitability in the dSub *in vivo*, which in turn support synaptic plasticity, neuronal oscillations and memory processing in mice. Considering the important role of the dSub in cognitive functioning in both animals and humans (Gabrielli et al., 1997; O’Mara et al., 2009; Small et al., 2000), our data indicate that Ca_V_3.1 T-channels may prove to be a promising novel drug target for cognitive deficits associated with multiple brain disorders.

## Acknowledgement

We thank the NeuroTechnology Center (NTC) of the University of Colorado Anschutz Medical Campus, for providing facilities to acquire video-EEG and behavioral data.

## Funding

Supported in part by funds from the Department of Anesthesiology at the University of Colorado Anschutz Medical campus, NIH grants K01 MH121567 (to S.M.J.), DP5 OD026407 (to J.A.H.), R01 HD097990, R01 GM118197, and CU Medicine Endowment (to V.J.-T.), R01 GM123746-03 and R35 GM141802-01 (to S.M.T.).

## Authors’ contributions

Performed miniscope *in vivo* calcium imaging experiments and analyzed the data: S.M.J., M.S.G., J.A.H. Performed LFP experiments and analyzed the data: S.M.J., R.V., J.A.H. Performed behavioral experiments and analyzed the data: S.M.J., N.B. Performed *ex vivo* electrophysiological experiments and analyzed the data: S.M.J., J.E.O. Performed IHC experiments: V.T., B.F. Designed the studies, supervised the overall project, and performed final manuscript preparation: S.M.J., J.A.H., V.J.-T., Y.R., P.S.H., and S.M.T.

---

**Figure 1–figure supplement 1.**
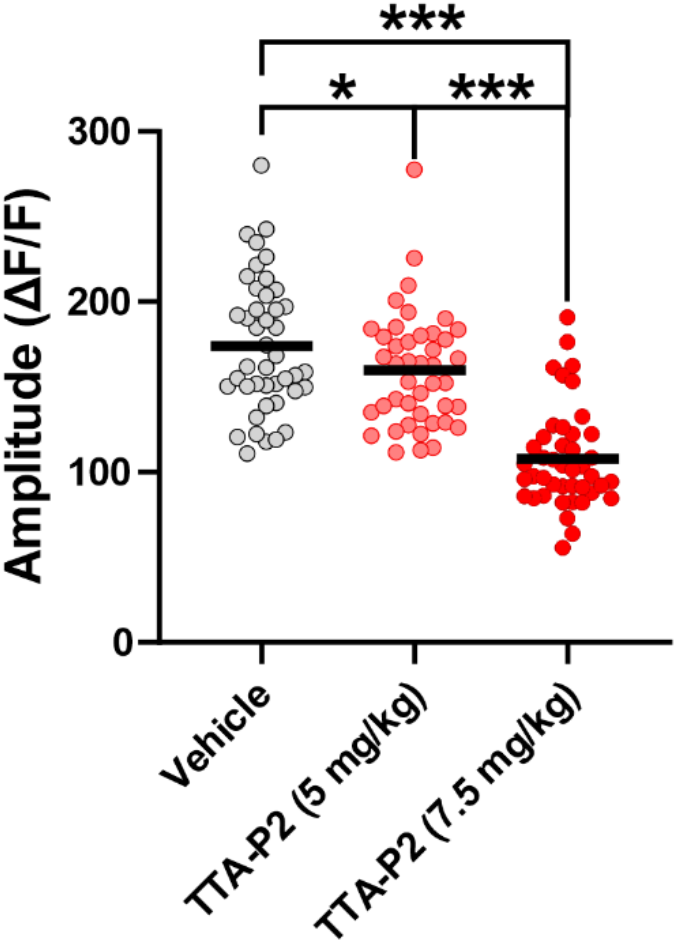
The effects of selective T-channel antagonist (TTA-P2) on dSub neuronal activity recorded using *in vivo* calcium imaging. The average amplitude of calcium transients recorded over three different sessions in the same population of dSub neurons (n=43): vehicle (gray), 5 mg/kg TTA-P2 (pink) and 7.5 mg/kg TTA-P2 (red). *P<0.05 and ***P<0.001, one-way RM ANOVA followed by Tukey’s *post hoc* test.

**Figure 3–figure supplement 1.**
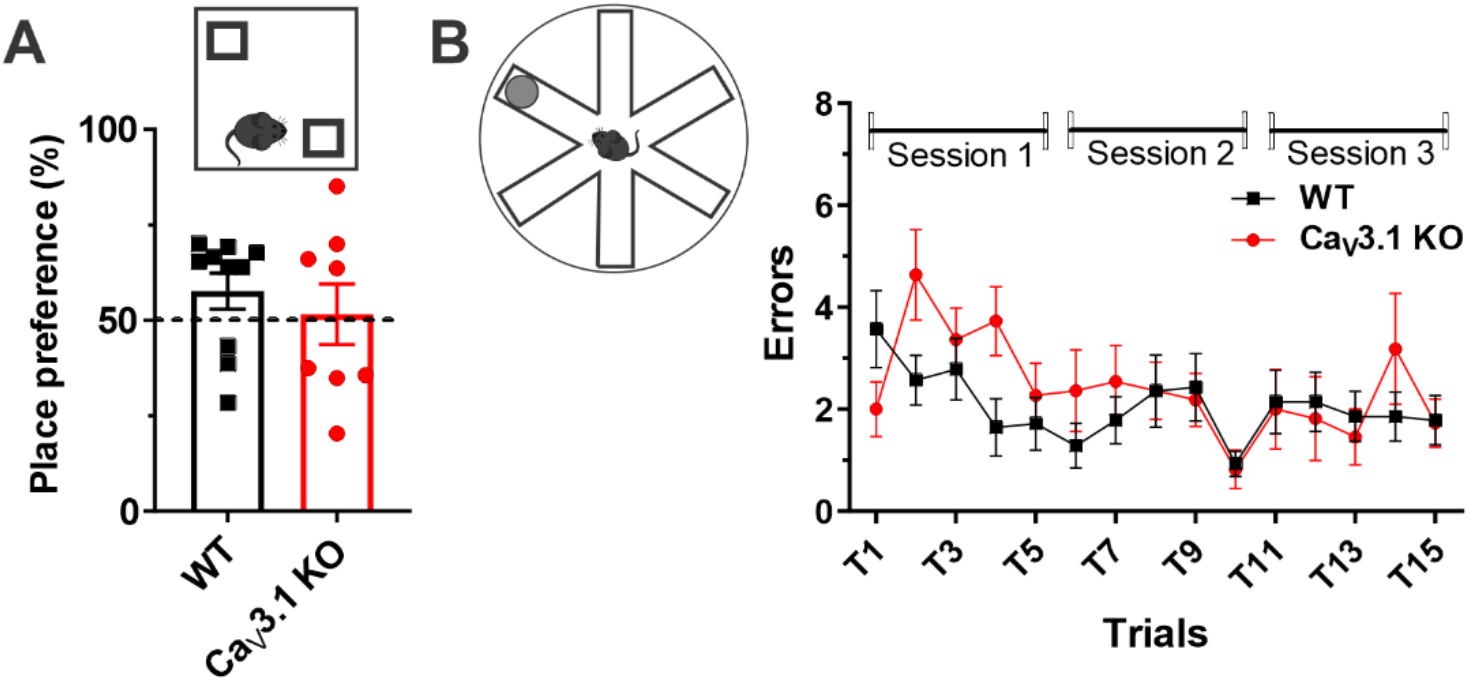
Behavioral assessment of cognitive functions in Ca_V_3.1 KO mice. **A.** The preference in the time exploring a displaced object in WT (n=10) and Ca_V_3.1 KO (n=8) mice during the novel place recognition test. **B.** The number of errors made by WT (n=14) and Ca_V_3.1 KO (n=11) mice during three swimming sessions in the radial arm water maze.

**Figure 5–figure supplement 1.**
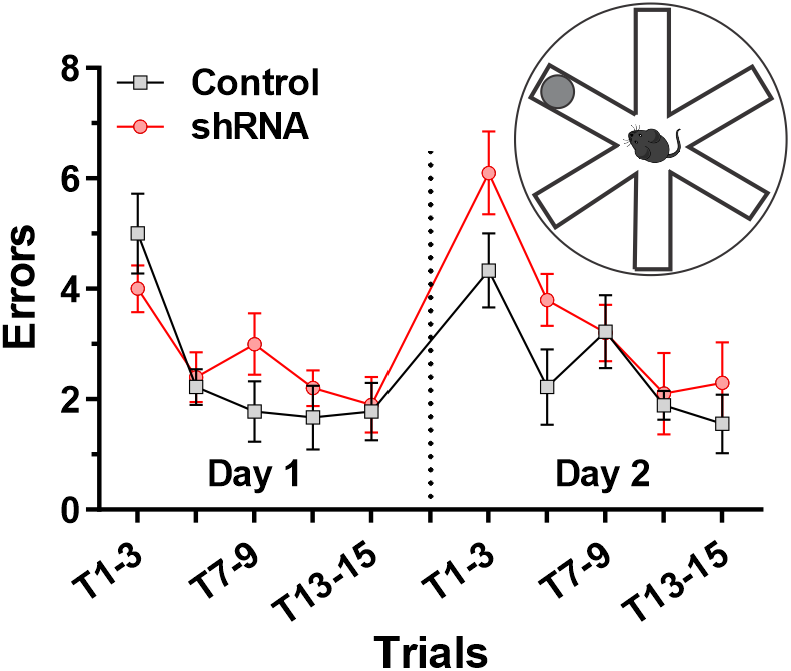
Behavioral assessment of cognitive functions in dSub-specific Ca_V_3.1 KD mice using the radial arm water maze (RAWM). The number of errors made by Control (n=9) and Ca_V_3.1-shRNA (n=10) mice during two days in the RAWM.

## Notes

### Competing Interest Statement

The authors have declared no competing interest.

